# Dengue Virus Surveillance in Nepal Yields the First On-Site Whole Genome Sequences of Isolates from the 2022 Outbreak

**DOI:** 10.1101/2024.06.02.597008

**Authors:** Rajindra Napit, Annie Elong Ngono, Kathie A. Mihindukulasuriya, Aunji Pradhan, Binod Khadka, Smita Shrestha, Lindsay Droit, Anne Paredes, Lata Karki, Ravindra Khatiwada, Mamata Tamang, Bimal Sharma Chalise, Manisha Rawal, Bimlesh Jha, David Wang, Scott A. Handley, Sujan Shresta, Krishna Das Manandhar

## Abstract

**Background:** The 4 serotypes of dengue virus (DENV1-4) can each cause potentially deadly dengue disease, and are spreading globally from tropical and subtropical areas to more temperate ones. Nepal provides a microcosm of this global phenomenon, having met each of these grim benchmarks. To better understand DENV transmission dynamics and spread into new areas, we chose to study dengue in Nepal and, in so doing, to build the onsite infrastructure needed to manage future, larger studies.

**Methods and Results:** During the 2022 dengue season, we enrolled 384 patients presenting at a hospital in Kathmandu with dengue-like symptoms; 79% of the study participants had active or recent DENV infection (NS1 antigen and IgM). To identify circulating serotypes, we screened serum from 50 of the NS1^+^ participants by RT-PCR and identified DENV1, 2, and 3 – with DENV1 and 3 codominant. We also performed whole-genome sequencing of DENV, for the first time in Nepal, using our new on-site capacity. Sequencing analysis demonstrated the DENV1 and 3 genomes clustered with sequences reported from India in 2019, and the DENV2 genome clustered with a sequence reported from China in 2018.

**Conclusion:** These findings highlight DENV’s geographic expansion from neighboring countries, identify China and India as the likely origin of the 2022 DENV cases in Nepal, and demonstrate the feasibility of building onsite capacity for more rapid genomic surveillance of circulating DENV. These ongoing efforts promise to protect populations in Nepal and beyond by informing the development and deployment of DENV drugs and vaccines in real time.

## Introduction

DENV, a member of the *Flaviviridae* family of single-stranded RNA viruses, exists as 4 serotypes and 19 genotypes and is mainly transmitted by the bite of *Aedes* mosquitoes.[1] About 20% of DENV infections lead to symptoms of dengue, ranging from self-limiting fever to hemorrhagic fever that can result in death.[2] Globally, the incidence of dengue has increased 30-fold over the past 50 years, and approximately 100 million cases are diagnosed annually.[3] The disease is endemic in more than 129 countries, with the majority of cases reported in Asia, South America, and the Western Pacific.[4–6] In Africa, there has been an upswing in dengue cases and outbreaks in recent decades[7, 8] and, in Europe and parts of the US, the presence of *Aedes albopictus* and climate change highlight the increasing threat of dengue to new regions. For example, in the US, the tropical/subtropical state of Florida experienced a record high number of DENV infections (966 travel-associated and 71 locally-acquired) during the 2022-2023 season.[9]

The outcome of DENV infection is influenced by both immune response history and viral genetics. Infection with DENV confers long-term protection against the same serotype, but only short-term protection against different serotypes; in fact, secondary infection with a different serotype can elicit severe dengue.[10, 11] Thus, a complex interplay between pre-existing antibody and T cell responses determines DENV disease severity.[11] DENV genotypes also play a critical role in modulating DENV pathogenesis, as well as transmission, based on associations between specific DENV genotypes and disease severity and epidemic intensity.[12–15] Informing control efforts (eg, vaccine design, testing, and implementation), requires regional and country-specific data on the precise genetics of circulating DENV strains and host immune status. Unfortunately, these data are lacking, due, in large part, to limited on-site infrastructure.

Nepal is a landlocked nation that spans 3 distinct topographic regions: a lowland tropical region bordering India, a country with a high dengue burden and high numbers of severe dengue cases even during primary DENV infections;[16] a temperate mountainous region bordering China, the most populous country in the world; and a subtropical/temperate elevated region in between. DENV was first detected in Nepal in 2004[17] and its distribution, while originally concentrated in the tropical region, has since extended to subtropical and temperate regions.[18–21] In Nepal, dengue outbreaks occur every year, with surges every 3 years since 2010 (ie, 2010, 2013, 2016, 2019, and 2022). In these surge years, the number of cases has increased from roughly 1500 in 2016 to 55,000 in 2022, and the dominant circulating serotype has alternated between DENV1 and 2: DENV1 in 2010[22] and 2016,[23] and DENV2 in 2013[24] and 2019.[21] There have also been shifts in genotype; for example, DENV2 IVb in 2004,[25] and Cosmopolitan IVa and Asian II in 2013.[22, 26] Similar DENV serotype and genotype dynamics have been recorded in India[27, 28] and China.[29, 30] Thus, Nepal offers a unique opportunity to investigate DENV evolution and transmission dynamics, both nationally and regionally involving 2 most populous countries in the world.

Before 2022, there were only a handful of studies on genetic diversity of DENV in Nepal[20, 31–34] – with only 1 study providing full genome sequences of 2 DENV2 isolates from 2015,[32] and the rest sequencing only the envelope (E) gene. To obtain sequencing data for DENV strains circulating in Nepal, we enrolled 384 patients with suspected dengue who visited Kathmandu’s Sukraraj Tropical and Infectious Disease Hospital during the 2022 outbreak. From these subjects, clinical and demographic data was collected, as well as 384 consented blood samples. After validating DENV infection (anti-DENV IgM, IgG, and DENV NS1 antigen), 50 of the 183 dengue-confirmed samples were analyzed for serotype, and near-complete genome sequences, of sufficient quality, obtained for 6. Our serotyping data confirmed that DENV1, 2, and 3 were mainly responsible for the 2022 outbreak.[35] Further, the 4 DENV1 and 3 sequences were closely related to strains circulating in India in 2019, suggesting they spread from India to Nepal via the shared border. In contrast, the 2 DENV2 sequences clustered with a 2019 strain from China, suggesting they spread from China, likely via air travel.

Most of this work was performed in Nepal, as a result of a 7-year-long effort to build the necessary intellectual and technical infrastructure. Indeed, ours is the first study to perform whole-genome sequencing of DENV in Nepal and, therefore, to demonstrate the feasibility of real-time genomic surveillance of DENV in this model country. As Nepal is a major tourist destination, this on-site capacity promises to play an important role in managing DENV spread within and beyond its borders.

## Results

All of the protocols and assays in this study were carried out in Nepal, except for the bioinformatics analyses, which were carried out in both Nepal and the United States (**Figure 1a**).

**Figure 1.**
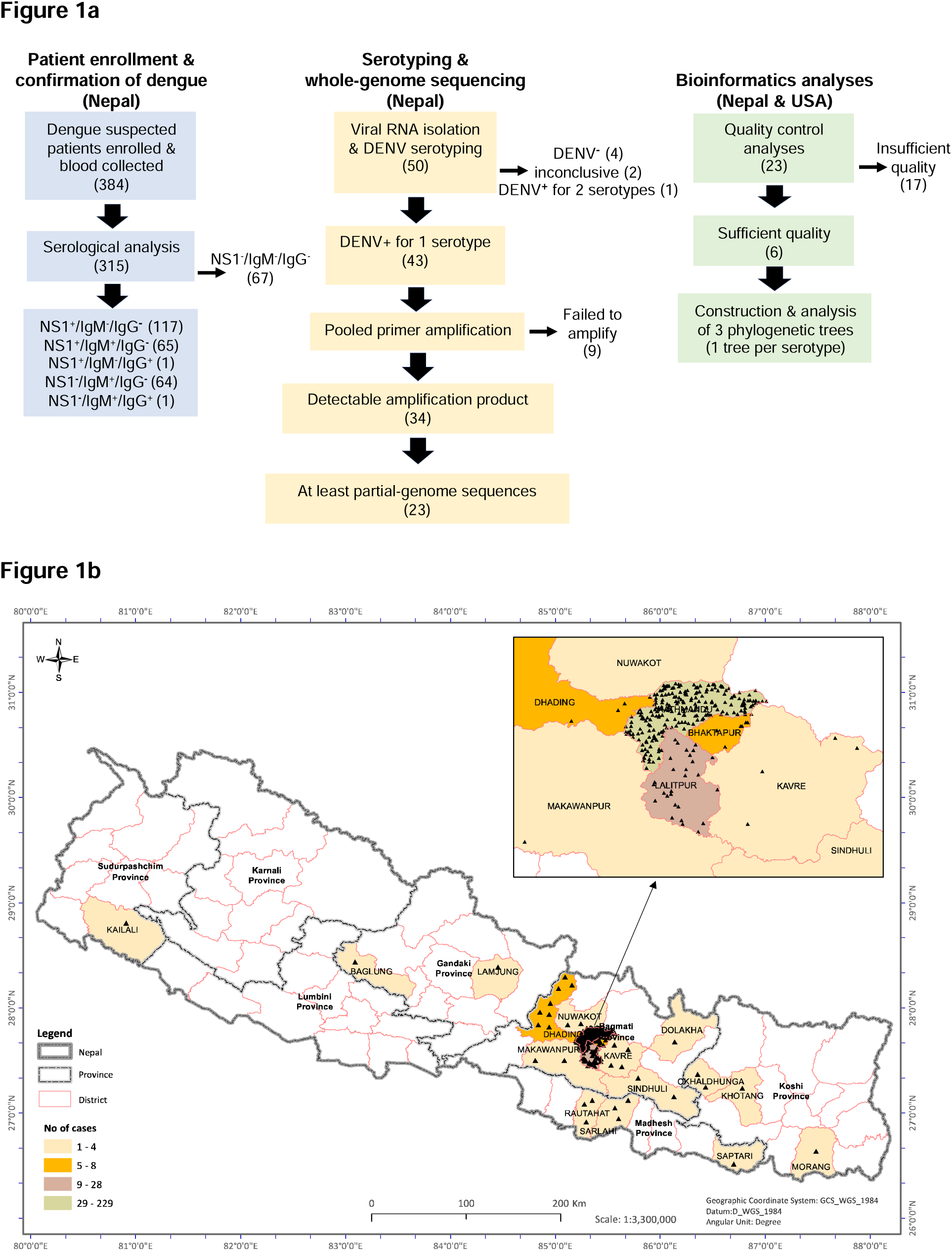
**(a) Project overview** (n values in parentheses). At Sukraraj Tropical and Infectious Disease Hospital in Nepal, 384 dengue-suspected patients were enrolled, blood samples collected, and NS1^+^ (dengue-confirmed) samples isolated (light blue boxes). At Tribhuvan University, also in Nepal, 50 samples were serotyped and whole-genome sequences obtained for 23 of them by NGS (yellow boxes). Finally, bioinformatic analyses were performed at Tribhuvan University in Nepal and Washington University/St Louis in the USA (green boxes). **(b) The vast majority of study participants were from temperate districts.** Demographic distribution of 296 of the 384 dengue-suspected cases. Enlargement, part of Bagmati province showing the highest density and number of dengue-suspected cases in Kathmandu and Lalitpur districts.

### Patient enrollment and confirmation of dengue

We enrolled 384 patients who presented with dengue-like symptoms at Sukraraj Tropical and Infectious Disease Hospital from September through December 2022. For each of the 384 study participants, a detailed clinical report was taken (**Table 1**), and a single blood sample collected. The most common presenting symptoms were fever and headache (81% and 70%, respectively), followed by muscle/joint pain (57%) and nausea (53%). Of the 312 febrile patients, 185 (59%) had intermittent fever, 120 (39%) had continuous fever, and 7 (2%) had remittent fever. Hemorrhage and/or rash, symptoms of severe dengue, were seen in 24 (6%) of study participants. Data for the time between fever onset and presentation at the hospital was available for 292 of the 384 participants (76%); 80% of the samples (235/292) were collected within 5 days following fever onset. Of the 384 samples collected, 328 were subjected to standard clinical blood cell testing; 134 (41%) had low platelet counts and 30 (9%) had low hemoglobin, signs of thrombocytopenia and anemia, respectively, which are both associated with dengue disease.[36, 37]

**Table 1.** Study population: demographic and clinical data.

| **Table 1. Study population: demographic and clinical data** |  |  |  |
| --- | --- | --- | --- |
|  | **Total** | **Male** | **Female** |
| **Age (years; n=384)** |  |  |  |
| <20 | 66 (17%) | 37 | 29 |
| 21-30 | 107 (28%) | 62 | 45 |
| 31-40 | 90 (23%) | 54 | 36 |
| 41-50 | 64 (17%) | 35 | 29 |
| 51-60 | 29 (8%) | 21 | 8 |
| >60 | 28 (7%) | 17 | 11 |
| **Residence (n=296, 77%)** |  |  |  |
| **Higher altitude temperate districts** | 287 (97%) | 173 | 114 |
| Kathmandu | 229 |  |  |
| Lalitpur | 28 |  |  |
| 22 other districts | 30 |  |  |
| **Lowland tropical districts** | 9 (3%) | 5 | 4 |
| Sarlahi | 3 |  |  |
| Rautahat | 3 |  |  |
| 3 other districts | 3 |  |  |
| **Symptoms (n=384)** |  |  |  |
| fever | 312 (81%) | 181 | 131 |
| headache | 271 (71%) | 153 | 118 |
| muscle/joint pain | 221 (58%) | 122 | 99 |
| nausea | 204 (53%) | 100 | 104 |
| lethargy | 172 (45%) | 92 | 80 |
| abdominal pain | 160 (42%) | 97 | 63 |
| vomiting | 98 (26%) | 47 | 51 |
| rash | 24 (6%) | 14 | 11 |
| hemorrhage | 24 (6%) | 14 | 10 |
| **Fever, pattern (n=312, 81%)** |  |  |  |
| continuous | 120 (39%) | 65 | 55 |
| intermittent | 185 (59%) | 112 | 73 |
| remittent | 7 (2%) | 4 | 3 |
| **Fever, days since onset (n=292, 76%)** |  |  |  |
| <6 | 235 (80%) | 148 | 87 |
| 6-12 | 40 (14%) | 18 | 22 |
| >12 | 17 (6%) | 9 | 8 |
| **Laboratory tests (n=328, 85%)** |  |  |  |
| **Platelets/μL blood** |  |  |  |
| <20,000 | 3 (1%) | 1 | 2 |
| 20,000-40,000 | 3 (1%) | 3 | 0 |
| 41,000-150,000 | 128 (39%) | 88 | 40 |
| >150,000 | 194 (59%) | 110 | 84 |
| **Hemoglobin ($\leq 12$ g/dL)** | 30 (9%) | 7 | 23 |
| **DENV serology (n=315, 82%)** |  |  |  |
| NS1 antigen only | 117 (37%) | 66 | 51 |
| IgM only | 64 (20%) | 40 | 24 |
| IgG only | 0 | 0 | 0 |
| NS1 & IgM | 65 (21%) | 41 | 24 |
| NS1 & IgG | 1 (<1%) | 1 | 0 |
| IgM & IgG | 1 (<1%) | 1 | 0 |
| Negative | 67 (21%) | 41 | 26 |
| **DENV serotype (n=50, 13%)** |  |  |  |
| DENV1 | 21 (42%) | 13 | 8 |
| DENV2 | 5 (10%) | 1 | 4 |
| DENV3 | 17 (34%) | 10 | 7 |
| DENV4 | 0 | 0 | 0 |
| DENV1 & 3 | 1 (2%) | 1 | 0 |
| Negative | 4 (8%) | 2 | 2 |

At the time of enrollment, the 384 study participants were residing in 5 of the 7 provinces of Nepal (**Figure 1b**). Precise residential location data were available for 296 participants (77%); of these, nearly all (287/296, 97%) were from higher altitude temperate districts, predominantly Bagmati province (altitude >1300 m) which includes Kathmandu (229/296, 77%), the most densely populated district. Only 9 study participants (3%) were from tropical lowland provinces (**Figure 1b**, **Table 1**). The ratio of males to females was 1.4 (59% male, 41% female), ages ranged from 9 to 82 years, with a median of 32, and 51% (197/384) were in the 21-to-40-year range (**Table 1**).

To screen for evidence of active or past DENV infections, serum from 315 participants was assayed for levels of anti-DENV IgM and IgG, and DENV NS1 antigen (**Table 1**). Of the 315 serum samples, 248 (79%) represented active dengue cases, based on the presence of DENV NS1 and/or anti-DENV IgM. Interestingly, 67 study participants (21%) were triple negative, despite presenting with dengue-like symptoms.

### Serotyping and whole-genome sequencing

Of the 117 serum samples that were positive for DENV NS1 antigen alone (**Table 1**, **Figure 1a**), 50, collected at the beginning of the outbreak, were subjected to RT-PCR to determine DENV serotype. Forty-four of the 50 samples (88%) were positive for 1 or 2 serotypes: DENV1 (21), DENV2 (5), DENV3 (17), and DENV1+3 (1); no samples tested positive for DENV4, 4 tested negative for all 4 serotypes, and 2 had no amplification of either sample or control RNA. Thus, during the 2022 outbreak in Nepal, DENV1 and DENV3 appear to have been the co-dominant circulating serotypes.

We next attempted to amplify the DENV sequences using pooled, serotype-specific tiled PCR primers, and 34 out of 43 samples yielded observable amplification product by gel electropheresis. The 9 samples that failed to amplify presumably had poor RNA quality or low viral genome abundance. Following library construction, NGS sequencing and bioinformatic analysis, we obtained at least partial DENV genome sequences from 23 samples (**Figure 1a**).

### Bioinformatics analyses

The sequencing data for the 23 DENV1, 2, and 3 isolates were subjected to quality control analyses to define near-complete genomes (>70% coding sequence coverage) for further analysis. Six of the 23 genomes satisfied this criterium: 3 DENV1, 2 DENV2, and 1 DENV3 (**Figure 1a, Table S1**). We used a custom database[38] to generate a maximum likelihood tree. All 3 DENV1 genomes clustered with genotype-V, suggesting that the circulating dominant DENV1 strains of the 2022 outbreak belong to genotype-V. The DENV2 genomes both clustered with Cosmopolitan-IVa genotype, and the DENV3 genome clustered with genotype-III (**Table S1**).

To predict the evolutionary relationships for our 6 DENV sequences, we first downloaded all DENV1-3 complete genome sequences submitted to the NCBI database after 2010; this yielded a custom database of 3429 DENV sequences: 2625 DENV1, 613 DENV2, and 191 DENV3. The database includes 3 DENV1 and 3 DENV3 genomes from the 2022 outbreak in Nepal that were sequenced in Israel.[34] We then constructed 3 phylogenetic trees (1 per serotype), using IQTree2, followed by timed ancestral analysis with TreeTime (**Figures 2-4**). Our 3 DENV1 genomes and the 3 genomes sequenced in Israel all shared a common ancestor dating back to 2019, and clustered closely with a 2019 DENV1 strain from India (MN923086); the next closest relatives were 2016 DENV1 strains from Singapore (MF033256, MF033261) and India (MK588396) – all with a 2013 common node (**Figure 2**). Our 2 DENV2 genomes clustered with DENV2 sequences from China, with the closest one reported in 2018 (MK564485); the common ancestral sequence of these 3 genomes most likely dates back to 2018 (**Figure 3**). Our 1 sequenced DENV3 genome and the 3 Israeli-sequenced DENV3 genomes clustered most closely with 2019 strains from India (ON123658, MK858155), and more distantly with numerous 2016-2019 sequences from China (**Figure 4**).

**Figure 2.**
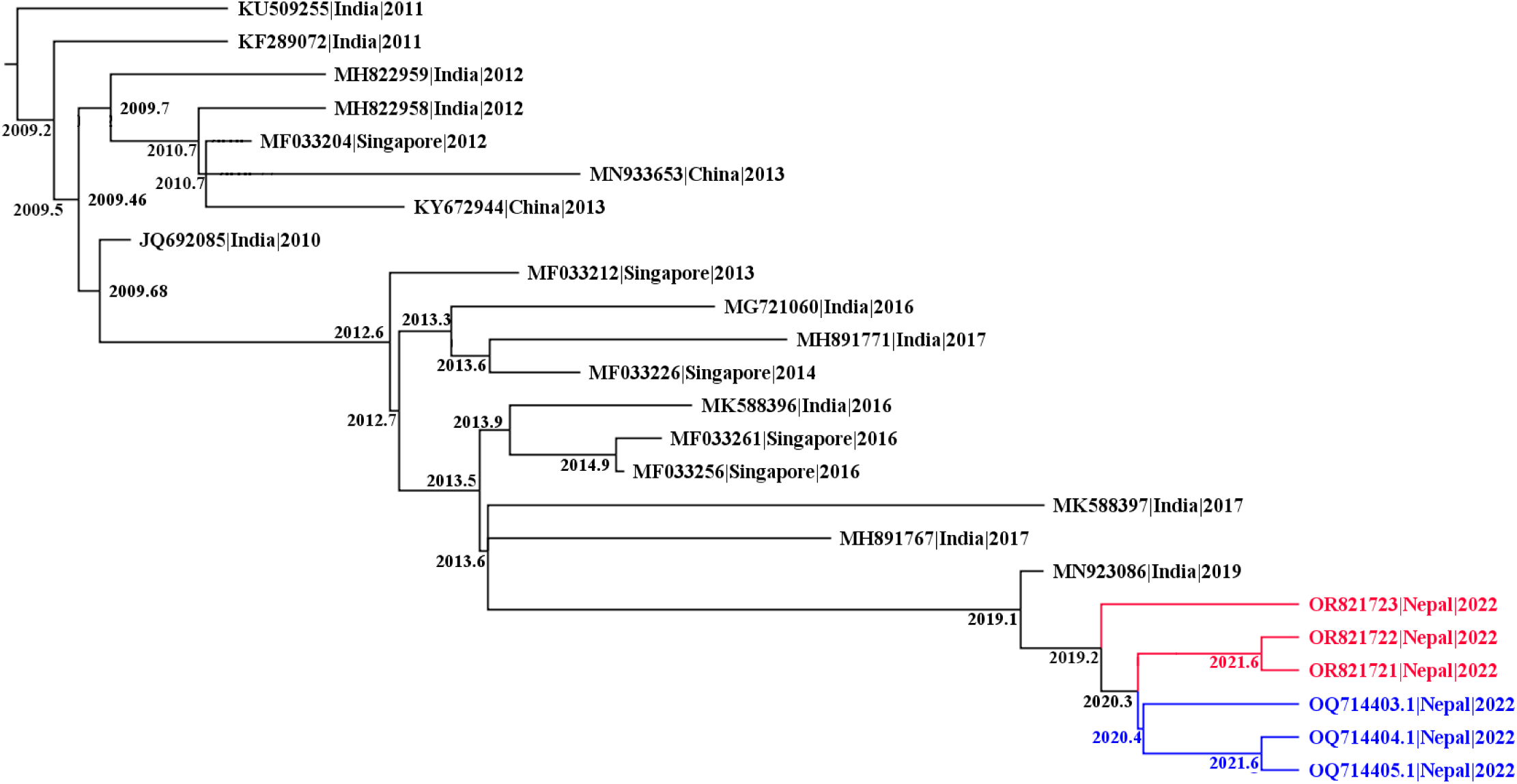
Phylogenetic tree of DENV1 full-length genomes. A maximum likelihood tree of DENV1 created using IQ-tree2, and dated to infer ancestral sequences with TreeTime. Sequences were named by accession number, country of origin, and year, and nodes labelled with the predicted year of the common ancestor; unrelated clades were collapsed. Red, 3 DENV1 genomes from this study (Table S1). Blue, 3 recently reported DENV1 genomes.

**Figure 3.**
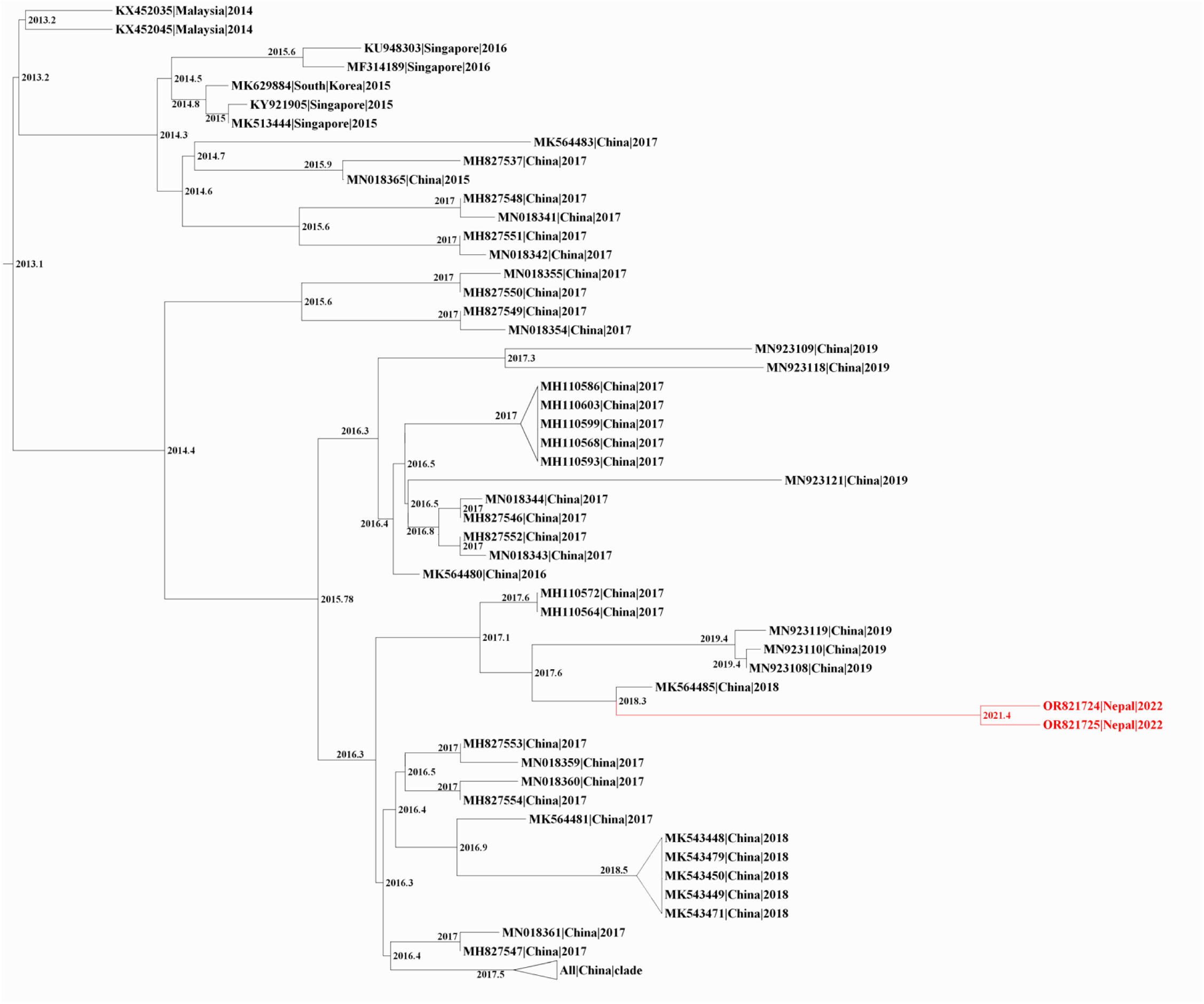
Phylogenetic tree of DENV2 full-length genomes. A maximum likelihood tree of DENV2 created using IQ-tree2, and dated to infer ancestral sequences with TreeTime. Sequences were named by accession number, country of origin, and date, and nodes labelled with the predicted date of the common ancestor; unrelated clades were collapsed. Red, 2 DENV2 genomes from this study (Table S1).

**Figure 4.**
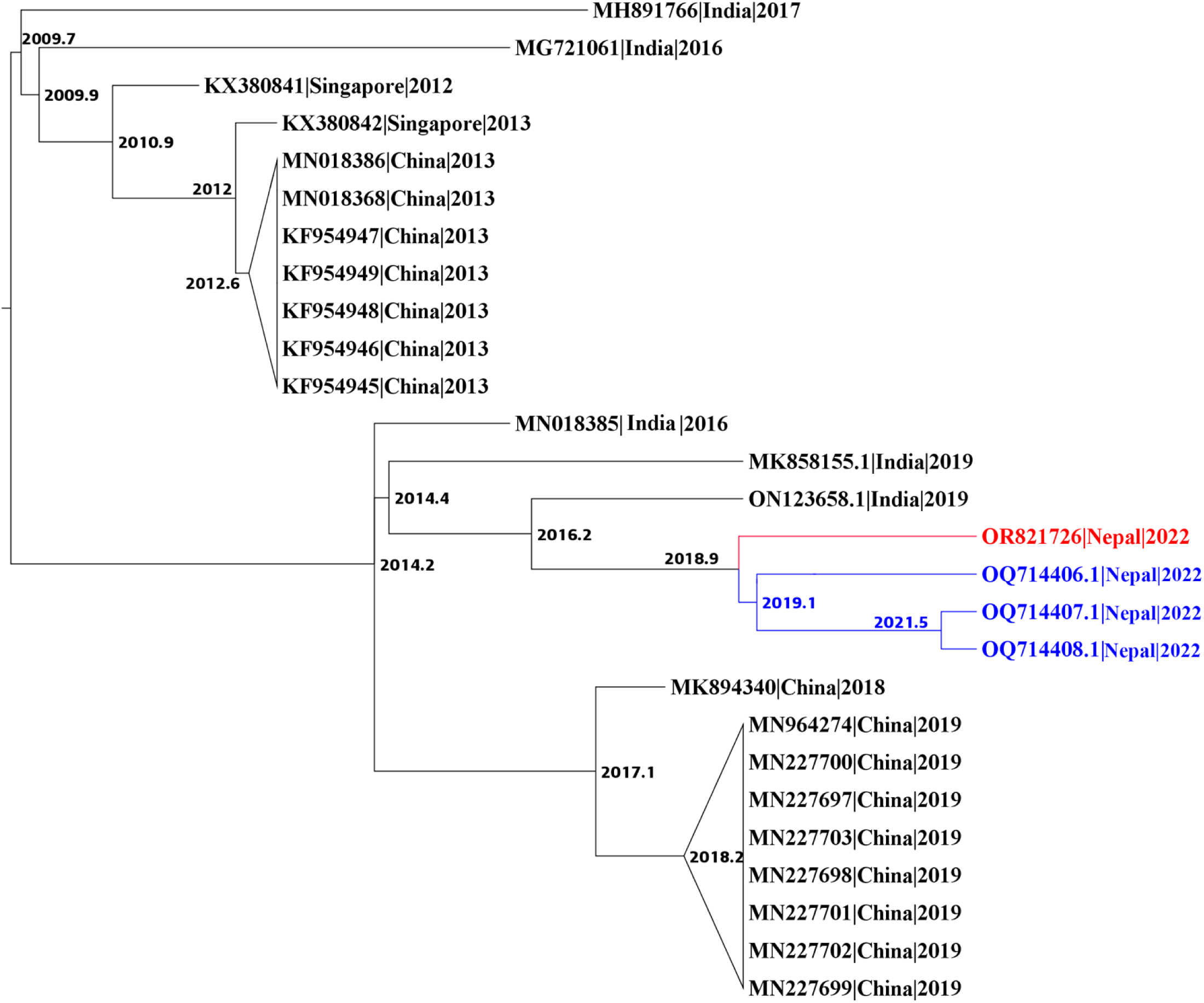
Phylogenetic tree of DENV3 full-length genomes. A maximum likelihood tree of DENV3 created using IQ-tree2 and dated to infer ancestral sequences with TreeTime. Sequences were named by accession number, country of origin, and year, and nodes labelled with the predicted year of the common ancestor; unrelated clades were collapsed. Red, 1 DENV3 genome from this study (Table S1). Blue, 3 recently reported DENV3 genomes.

To better understand the evolution of our 2022 Nepalese DENV genomes, it was important to compare them to strains in Nepal prior to 2022. Unfortunately, there are no available full genome sequences of DENV1 and 3 from dengue outbreaks in Nepal prior to 2022, and only 2 full sequences for DENV2 (sequenced by our group)[32]. We therefore compared E gene sequences of our 5 DENV1 and 2 genomes against other DENV1 and 2 sequences from Nepal in NCBI GenBank, and their respective closest sequences (eg, for DENV1, MN923086 and MF033256 from India and Singapore, respectively); E-gene sequences for DENV3 strains from Nepal are not available. This analysis revealed that our 3 DENV1 genomes are more closely related to a 2019 strain from India, than to 2010 and 2017 DENV1 strains from Nepal (**Figure S1**), and our 2 DENV2 genomes are more closely related to 2018 and 2019 strains from China than strains reported from pre-2022 outbreaks in Nepal (**Figure S2**). Taken together, our complete genome and E gene sequence analyses suggest that the 2022 dengue outbreak in Nepal was driven by recent regional travel and importation from India and China.

To determine the extent and types of missense mutations in our 6 DENV1-3 genomes, we used our custom NCBI-derived database to compare the complete sequences of our 6 strains against their respective reference sequences (**Figure S3, Table S2**). For our 3 DENV1 strains, we discovered 6 distinct missense mutations in E and NS5 genes, 2 of which were shared by all 3 strains. Analysis of our 2 DENV2 strains revealed 7 distinct missense mutations in 6 genes (capsid, E, and 4 NS); 3 of these mutations were present in both strains. Our DENV3 strain had 8 distinct missense mutations in 6 genes (E, 2 K, and 4 NS). Thus, E and NS5 genes appear to be mutational hotspots in our 6 DENV genomes.

## Discussion

Dengue cases are increasing globally, especially in endemic countries in South America and Southeast Asia – including Nepal, where cases in the 2022 outbreak were 3-fold higher than 2019. DENV infection is also spreading into new geographic areas, including Europe.[6] In the US, locally-acquired dengue cases have been reported in Texas, Arizona, Hawaii, and California and, in Florida, over 1000 DENV infections (71 locally acquired) were reported during the 2022-2023 dengue season.[9] Addressing this public health threat will require scalable real-time DENV genomic surveillance programs. This study is part of an ongoing effort to build such a program.

Some of the global spread of dengue has been from tropical and subtropical regions to temperate ones. In Nepal, for all reported cases, 3% were from the temperate region in 2016, and 50% in 2022. In our current study, participants were from 5 of the 7 provinces of Nepal, with most living in higher-elevation temperate districts. This is consistent with a 2023 report from Nepal’s Ministry of Health and Population showing, for the first time, dengue cases in all 7 provinces and 77 districts, and that higher-elevation temperate districts were most affected.[39] This spread to higher altitude temperate regions is likely due to a combination of DENV evolution, mosquito adaptability, and climate change.[40]

Of the 50 samples subjected to serotyping, 39 tested positive for DENV1 and/or 3, and only 5 tested positive for DENV2 (no samples tested positive for DENV4). These data suggest that during the 2022 outbreak in Nepal, DENV1 and DENV3 were the co-dominant circulating serotypes, and are consistent with other studies reporting circulation of multiple serotypes during the 2022 outbreak in Nepal.[34, 35, 41] This is the first dengue outbreak in Nepal with 3 serotypes co-circulating, 2 serotypes co-dominant, and DENV3 as 1 of the circulating serotypes. Thus, in less than 2 decades (ie, since the first reported dengue case in Nepal in 2004), Nepal is on a trajectory to become hyperendemic, with all 4 serotypes co-circulating by the next predicted outbreak in 2025. This is a cautionary tale for countries where dengue was recently introduced – such as France,[42] Italy,[43] and the US.[9]

From 23 samples, we generated 17 partial and 6 near-complete (>70% coverage) genomes, using a published direct sequencing strategy with tiled primer amplicons.[44] With this approach, coverage can be reduced by lower sample quality, lower viral load, and higher point mutations.[45, 46] In addition, our primers were designed using 2016 consensus sequences in the genome database, and might, therefore, be susceptible to viral evolution leading to amplicon dropouts.[44, 47] This underscores the need for continual refinement of tiled primers for optimal amplification of circulating virus strains, as documented during the COVID pandemic by the need to update ARTIC tiled primers for sequencing SARSCoV-2 genomes.[47]

For the 6 near-complete DENV1-3 sequences in this study, the mutations were concentrated in E and NS5 genes (3 DENV1 isolates), and in E, NS2B, and NS3 genes (2 DENV2 isolates); in the single DENV3 isolate sequenced, mutations were distributed more evenly across the genome. E gene mutations are likely linked to immunological pressure[48] and NS5 mutations to selection pressure on virus in *A*.aegy*pti*.[49] Hence, mutations discovered by our whole genome sequence analysis could be linked to disease severity and virus transmission, and are important to consider when designing and testing vaccines and antivirals.

Our 3 DENV1 isolates are genotype V and are closely related to a strain responsible for a 2019 dengue outbreak in India that, in turn, is distantly related to 2016 isolates reported in Singapore, and India. This is consistent with a recently published analysis of 3 other DENV1 isolates from the 2022 outbreak.[34] Our 2 DENV2 isolates are cosmopolitan IVa genotype, consistent with previous reports for DENV2 in Nepal,[31, 33] and are closely related to a 2018 isolate from China[50] and isolates responsible for the 2017 outbreak in China. Finally, our single DENV3 isolate is genotype III; this isolate, along with 3 other DENV3 isolates from the 2022 outbreak in Nepal,[34] is closely related to an Indian strain isolated in 2019. Thus, our study and the one by Zuckerman et al. strongly suggest that the 2022 DENV1 and 3 Nepalese isolates were from India (DENV1 and 3), and our study suggests that the DENV2 isolates were from China.

Our findings point to regional transmission dynamics between Nepal, India – one of the high dengue burden countries and second most populous country, and China – the most populous country. While severe dengue is usually seen in secondary DENV infections, it was recently reported in India during primary infections,^16^ suggesting that DENV-host interactions may differ depending on the population, geographical location, and other environmental factors. This all points to the need for both local and larger regional surveillance mechanisms. Our multi-year international collaborative effort demonstrates the feasibility of building mutually beneficial on-site genomic surveillance infrastructure in low-income countries. The resulting research capacity in Nepal will have broad impact. It provides a unique opportunity to study DENV evolution and transmission in South and East Asia, while elevating the careers of researchers in both high- and low-income countries.

## Materials and Methods

### Ethics statement

This study was approved by the Nepal Health Research Council (NHRC) who forms the Ethical Review Board (ERB) (reg. 686/2021). The current version of the guideline followed by NHRC is based on the basic principles of the Nuremberg Code, the World Medical Association (WMA) Declaration of Helsinki, the Council of International Organization of Medical Sciences (CIOMS), International Ethical Guidelines for Biomedical Research Involving Human Subjects, the World Health Organization (WHO), International Conference on Harmonization (ICH) and Guidelines for Good Clinical Practice (GCP). All patients or their parent/guardian (if patient was <18 years of age) provided informed consent by signature and/or fingerprint after reading the consent form or having it read to them in their language.

### Study population, enrollment criteria, and sample collection

In this cross-sectional study, we enrolled 384 patients who visited Sukraraj Tropical and Infectious Disease Hospital (STIDH) in Kathmandu from September 2022 through December 2022 (ie, 4 months). Enrollment criteria were 2 or more of the following symptoms: temperature >38°C for more than 1 week, nausea, body/muscle or abdominal pain, vomiting, or rashes. Clinical and demographic data were recorded by the attending physician, and ArcMap 10.4 software (ArcGIS) used to represent the geospatial distribution of dengue cases. Single blood samples (3-5 mL) were collected at enrollment, and sera immediately isolated by centrifugation and stored at −20°C.

### NS1 antigen detection and anti-IgM/IgG DENV serology

Serum samples were analyzed for the presence of DENV NS1 antigen and anti-DENV IgM and IgG using Bioline Rapid Diagnostic Test kits (Abbott, 11DD104A), per manufacturer’s instructions. Samples with inconclusive results were re-tested; if a result remained inconclusive, the sample was considered negative.

### Viral RNA isolation and DENV serotyping

Viral RNA was extracted from 140 μl of serum using the spin protocol of the QIAamp Viral RNA Mini Kit (Qiagen, 52904), and DENV1-4 serotypes detected using a real time RT-PCR multiplex diagnosis kit (Center for Disease Control and Prevention, KK0129) that included positive control DENV 1-4 RNA, human specimen (positive) control (human ribonuclease P RNA; HSC), primers, and DENV1-4 serotype specific probes. Each reaction well contained 5 µL of RNA (extracted viral, DENV, or HSC) or negative control (RNase-free water) plus 20 µL of PCR reaction mix: 2.2 μL RNase-free water, 12.5 μL 2X Premix-buffer, 0.5 μL forward and reverse primers for DENV1 and DENV3 (2 μL total), and 0.25 μL forward and 0.25 μL reverse primers for DENV2 and DENV4 (1 μL total), 0.45 μL probes for each serotype (1.8 μL total), and 0.5 μL Superscript III Platinum One-Step qRT-PCR system enzyme (Invitrogen, 11732– 020). PCR was performed using a CFX96 Touch Real-Time PCR Detection System (Bio-Rad): reverse transcription (50°C, 30 min), inactivation (95°C, 2 min), followed by 45 cycles of 95°C for 15 s and 60°C for 1 min. DENV serotypes were identified using serotype-specific probe amplified with a maximum Ct value of 37: DENV1 (FAM/blue), DENV2 (Hex/green), DENV3 (Texas red), and DENV4 (Cy5/purple).

### Amplicon generation and validation for library preparation

Extracted viral (and control) RNAs were reverse transcribed using an iScript cDNA Synthesis Kit (Bio-Rad, 1708891), per manufacturer’s instructions. Previously designed serotype-specific primers for DENV1-3 (10 pairs per serotype) were validated individually by PCR of positive control DENV 1-3,[44] and validated primers balanced and divided into 2 pools per serotype (ie, 6 pools total; DENV4 was not detected in any of the studied samples). Amplicons were generated in 2 different 25 µL PCR reactions (9.5 µL nuclease-free water, 12.5 µL 2x QIAGEN Multiplex PCR Master Mix, 2 µL balanced primer pool 1 or 2 and 1 µL cDNA) using a T100^TM^ Thermal Cycler (Bio-Rad) and the following conditions: initial denaturation (94°C, 15 min) followed by 40 cycles of denaturation-annealing-extensions (denaturation [94°C, 30 s], annealing for 30 s [47°C, DENV1; 53°C, DENV2; 42°C, DENV3], and extensions [72°C, 3 min], followed by a final extension [72°C, 5 min]. Amplicon products were visualized in 1.5% agarose gels, cleaned using Ampure XP^TM^ magnetic beads, and quantified with the Qubit dsDNA HS Assay Kit (Thermo-Fisher, Q33230).

### Whole-genome sequencing

DNA libraries were prepared using tagmentation, where amplicons are subjected to fragmentation and adaptor ligation in a single step using Illumina’s DNA library prep kit (20051274). Each sample (and a no-DNA negative control) was given a unique barcode using IDT Illumina DNA PCR Indexes kit (20050258), per manufacturer’s instructions. Barcoded libraries were pooled in groups of 5 samples (34 samples and the no-DNA control in 7 pools; **Figure 1**). Each pool was quantified with the Qubit dsDNA HS Assay Kit, subjected to fragment analysis on the Agilent 4200 TapeStation using High Sensitivity DNA ScreenTape Analysis (Agilent), normalized to 4 nM, and then combined into a single pool. This single pool was then denatured using NaOH (0.2 N), diluted to 12 pmol, and sequenced on a MiSeq platform using Reagent Kit v3 (Illumina, MS-102-3003).

### Consensus sequence generation

Paired-end reads were processed through a multistage quality-control procedure to remove primers, adapters, and low-quality sequence data.[51] Individual sample assemblies were generated using MEGAHIT.[52, 53] Assembled contigs were assigned taxonomy by querying NCBI’s BLASTn with default parameters. The reference with the highest identity to the longest DENV contig in a sample was retained for reference-guided assembly, which was performed by mapping reads to the selected DENV reference genome using BWA-MEM with default parameters.[54] The aligned SAM file was then sorted via SAMtools,[55] and primary alignments extracted (samtools view -h -F 2048) and filtered to require a mapping quality ≥30 (samtools view -bq 30). The filtered BAM was sorted and read groups replaced with a single read group using Picard[56] to satisfy GATK input file requirements (picard SortSam SO=coordinate and picard AddOrReplaceReadGroups). Variants were called using GATK (gatk3 -T UnifiedGenotyper --genotyping_mode DISCOVERY -stand_call_conf 30 -ploidy 1 - defaultBaseQualities 60),[57] and consensus genome generated using vcf2genome (-minc 3 - minq 30 -minfreq 0.9).[58] Following reference-guided assembly, consensus genomes were compared a second time to the NCBI nt database using BLASTn with default parameters. If a DENV reference genome was identified with higher similarity to the new consensus genome, it was retained and variants and consensus genomes recalculated as above. Genome diagrams were generated in R with custom code using ggplot2 v3.4.2 from tidyverse v2.0.0.[59]

### Phylogenetic and statistical analyses

Genotyping checks were performed using the online DENV typing tool and an open-source database of DENV sequences, as previously described.[60] Sequences were aligned using MAFFT[61] and visualized in Aliview;[62] alignments were subjected to manual quality control for any insertion of ‘n’ against the reference sequences, and bad sequences removed. Maximum likelihood trees were created using IQ-TREE 2,[63] with auto selection for the nucleotide substitution model and 1,000 bootstraps for branch support. To determine the clusters/origin of each sequenced sample, another dataset was obtained from the NCBI database for all DENV1-3 sequences submitted after 2010, a maximum likelihood tree created with IQ-TREE 2, and timed with TreeTime.[64]

## Supporting information

Supplementary files. Napit et al

## Data sharing

All relevant data are included in the manuscript or supplemental data. Sequences are available in NCBI gene bank (accession ID-OR821721-OR821726).

## Ethical Approval statement

This study was approved by the ethics review boards of the Nepal Health Research Council (reg. 686/2021).

## Acknowledgements

We acknowledge the NIAID Center for Research in Emerging Infectious Diseases (CREID) Network, especially the CREID-ESP (Epidemiology, Surveillance, Pathogenesis) for its engagement and devotion to strengthening research capacity (genomic and laboratory practices) in Nepal. We thank the Handley and Wang Labs (Washington University School of Medicine St-Louis) for providing bioinformatic and NGS trainings at Tribhuvan University, Dr. Laurence Cagnon and the Shresta Lab (La Jolla Institute for Immunology, LJI) for providing trainings at Tribhuvan University on biosafety practice, REDCap, sample collection, and manuscript preparation and submission.

## Authors contributions

RN, AEN, KDM, and SS conceived the study. BSC, MR, RK, and BK enrolled patients and collected metadata. BK, SmS, LK, RK, MT, and BJ performed the experiments. RN and KDM supervised the experiments. RN, AEN, KAM, and AP analysed and interpreted the data. RN, AEN, KDM, and SS wrote the manuscript; RN, AEN, KDM, KAM, LD, AnP, SH, DW, and SS edited the manuscript drafts, and all authors approved the final version.

## Data availability statement

The data underlying this article are available in the article and its online supplementary material. Any supplementary data will be shared on reasonable request to the corresponding author.

