## Supplementary files. Napit et al for "Dengue Virus Surveillance in Nepal Yields the First On-Site Whole Genome Sequences of Isolates from the 2022 Outbreak"

**Figure S1**

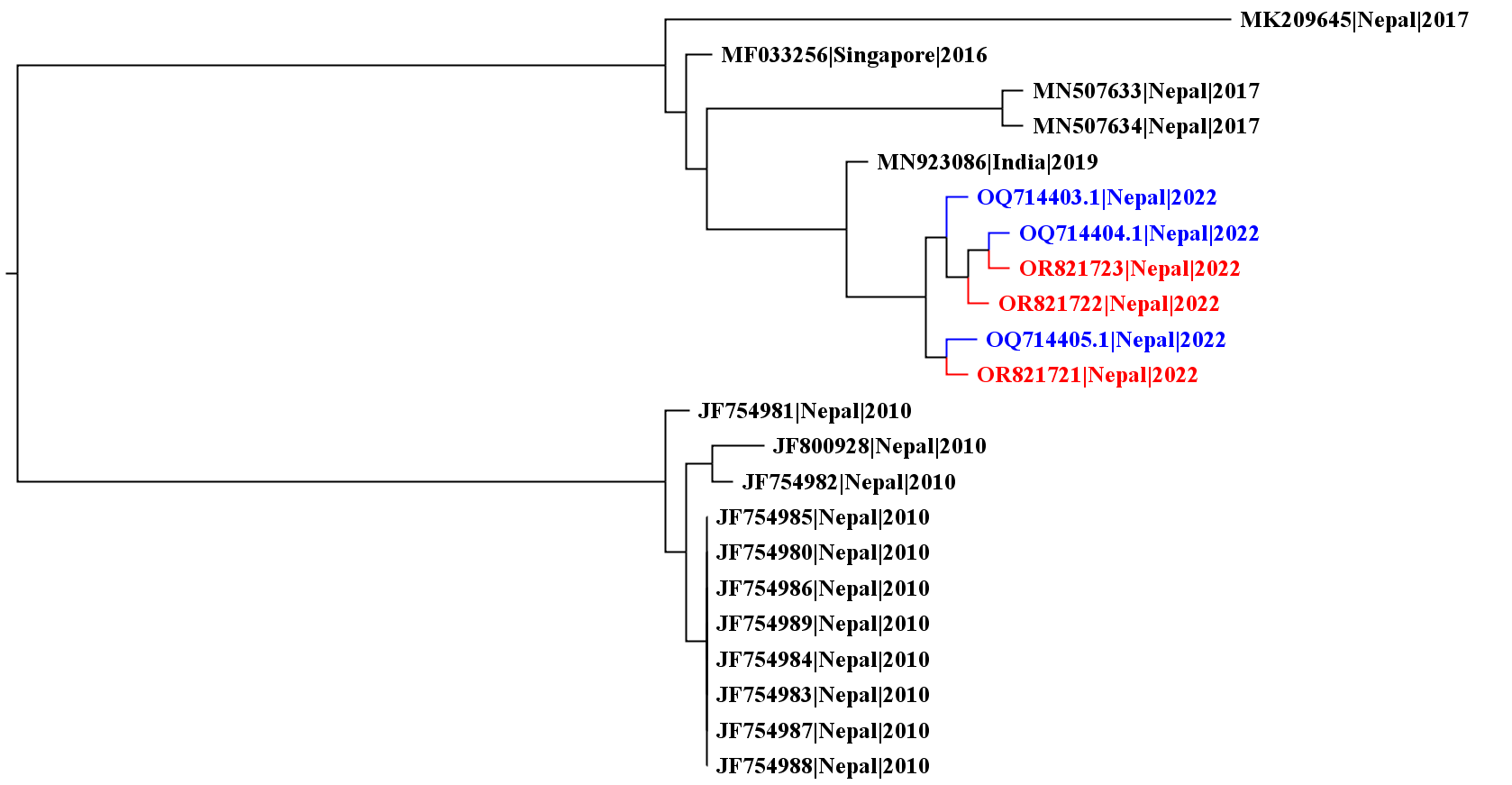

**Supplementary Figure 1. Comparison of E gene sequences of published DENV1 strains in Nepal (2010-2022).** E-gene-based maximum likelihood tree of DENV1 created with IQ-tree2, and dated to infer ancestral sequences with TreeTime. Sequences were named by accession number, country of origin, and year. Red, 3 DENV1 strains from this study (Table S1); Blue, 3 recently reported DENV1 strains.

**Figure S2**

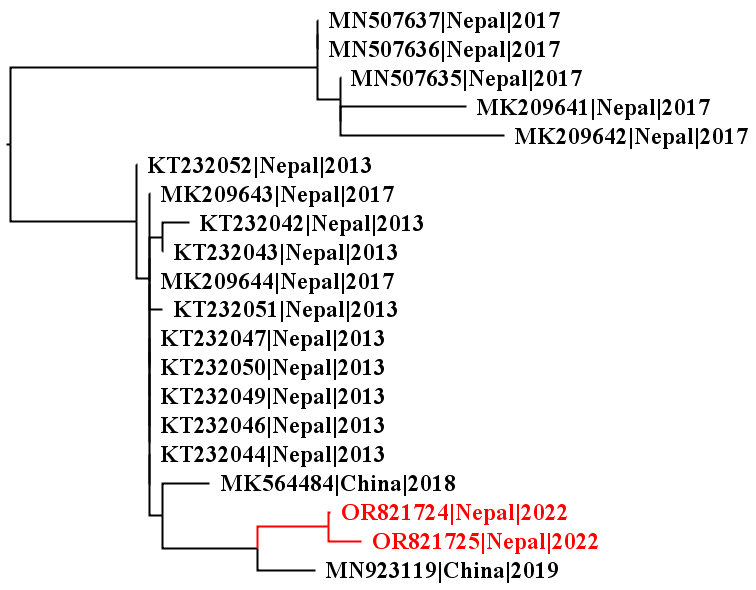

**Supplementary Figure 2. Comparison of E gene of published DENV 2 strains in Nepal (2013-2022).** E-gene-based maximum likelihood tree of DENV2 created with IQ-tree2, and dated to infer ancestral sequences with TreeTime. Sequences were named by accession number, country of origin, and year. Red, 2 DENV2 strains from this study (Table S1).

**Figure S3**

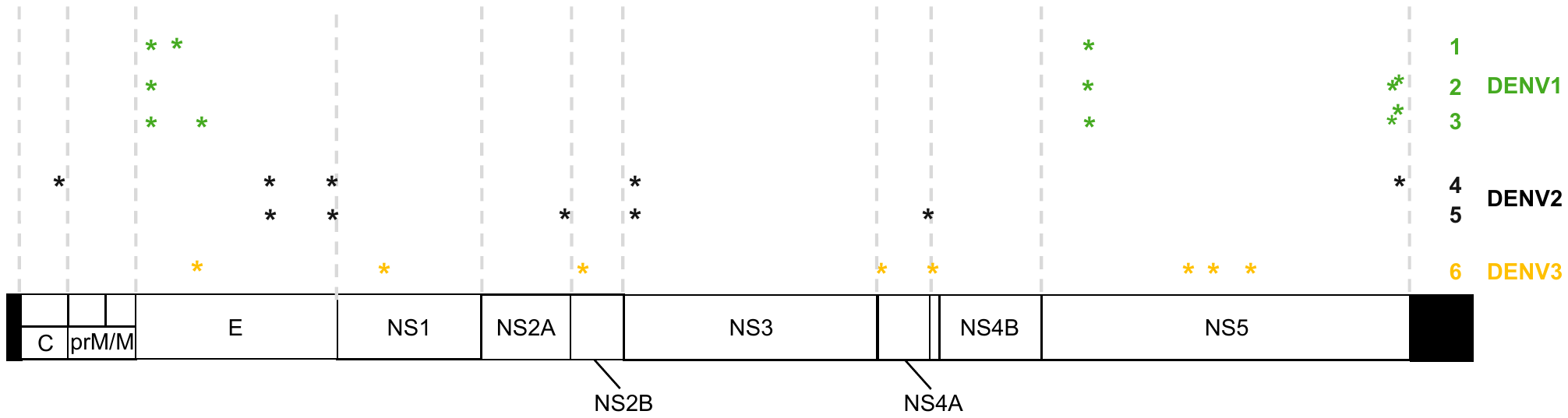

**Supplementary Figure 3. Genomic variation in 6 DENV strains.** Six genomes from this study – 3 DENV1, 2 DENV2, and 1 DENV3 (Table S1) were compared against their closest respective reference sequence in the NCBI-derived database (DENV1 vs MN923086.1, DENV2 vs MW512481.1, DENV3 vs MN018385.1) and missense mutations identified (asterisks). C, capsid; E, envelope; NS, nonstructural protein; prM/M, preMembrane/membrane.

| **Supplementary Table 1. Genotypes for DENV isolates** | | |
| --- | --- | --- |
| **Isolate** | **Serotype** | **Genotype** |
| 1 | 1 | V |
| 2 | 1 | V |
| 3 | 1 | V |
| 4 | 2 | Cosmopolitan IVa |
| 5 | 2 | Cosmopolitan IVa |
| 6 | 3 | III |

| **Supplementary Table 2. Nucleic acid changes and corresponding amino acid changes in DENV isolates vs reference sequences.** | | | | | |
| --- | --- | --- | --- | --- | --- |
| **Serotype** | **Isolate** | **Position** | **Feature** | **Base change** | **Amino acid change** |
| **DENV1** | **1** | 1084 | E | G > A | Arg > Lys |
|  |  | 1284 | E | A > T | Ile > Leu |
|  |  | 7881 | NS5 | C > T | Ser > Leu |
|  | **2** | 1084 | E | G > A | Arg > Lys |
|  |  | 7881 | NS5 | C > T | Ser > Leu |
|  |  | 10122 | NS5 | G > A | Val > Ile |
|  |  | 10155 | NS5 | T > G | Pro > Ala |
|  | **3** | 1084 | E | G > A | Arg > Lys |
|  |  | 1443 | E | T > C | Phe > Leu |
|  |  | 7881 | NS5 | C > T | Ser > Leu |
|  |  | 10122 | NS5 | G > A | Val > Ile |
|  |  | 10155 | NS5 | T > G | Pro > Ala |
| **DENV2** | **4** | 435 | ancC | G > A | Met > Ile |
|  |  | 1928 | E | C > T | Ser > Phe |
|  |  | 2386 | E | G > A | Val > Ile |
|  |  | 4565 | NS3 | G > A | Arg > Lys |
|  |  | 10201 | NS5 | G > A | Gly > Glu |
|  | **5** | 1928 | E | C > T | Ser > Phe |
|  |  | 2386 | E | G > A | Val > Ile |
|  |  | 4192 | NS2A | C > T | Leu > Phe |
|  |  | 4565 | NS3 | G > A | Arg > Lys |
|  |  | 6861 | NS4B | C > T | Thr > Ile |
| **DENV3** | **6** | 1413 | E | C > T | Ala > Val |
|  |  | 2796 | NS1 | C > T | Thr > Ile |
|  |  | 4241 | NS2B | A > G | Thr > Ala |
|  |  | 6438 | NS4A | A > G | Asn > Ser |
|  |  | 6810 | 2K | C > T | Thr > Ile |
|  |  | 8679 | NS5 | T > C | Val > Ala |
|  |  | 8867 | NS5 | G > A | Asp > Asn |
|  |  | 9133 | NS5 | T > G | Asn > Lys |
| ancC, anchored capsid; E, envelope; NS, nonstructural | | | | | |
